# Nutritional responses of bumblebees to thermal stress

**DOI:** 10.64898/2026.03.09.710642

**Authors:** Coline Monchanin, Stéphane Kraus, Jonathan Gerbore, Jean-Marc Devaud, Juliano Morimoto, Mathieu Lihoreau

## Abstract

Extreme climatic events impose considerable stress on organisms with consequences for key ecological interactions such as pollination. Because temperature directly affects metabolic processes, heat variations may also importantly influence the nutritional needs and feeding choices of animals. Here, we studied the effects of thermal stress on the nutritional choices and performances of bumblebees, using a 3D nutritional geometry design. At optimal temperature for colony development (30°C), bees successfully balanced carbohydrate, protein, and lipid collection, at levels beneficial for body weight and survival. Under cold stress (20°C), bees reduced their overall nutrient collection while selecting proportionally more carbohydrates, thereby prioritizing survival over weight gain. Under heat stress (35°C), nutrient balancing was disrupted and survival dropped. Notably however, across all temperatures, bees maintained stable lipid collection while flexibly adjusting the amount of carbohydrates and proteins, suggesting strong constraints on lipid regulation. Given the pivotal role of bees for pollination, identifying how their nutritional needs change in response to climatic conditions is of prior importance for food safety and the conservation of terrestrial ecosystems.

## Introduction

Understanding how animals adapt to rising ambient temperatures is a major challenge for predicting population declines [1], species distributions [2], and broader ecological consequences of climate change such as disruptions of essential ecosystem services [3]. At the most basic level, changes in temperature can influence both the nutritional needs of organisms and the composition of food resources available to them [4]. Yet, little is known about how, and to what extent, animals can adjust their feeding behaviour to maintain high performance across thermal gradients [5].

Animals, from mammals to insects, actively regulate their intake of multiple nutrients to optimise key fitness traits, often by foraging on complementary food sources [6]. This nutrient regulation typically involves trade-offs between the optimisation of fitness traits such as survival, reproduction, and development [7–10]. Temperature can shift these nutritional trade-offs by affecting food availability, nutrient composition, and the energetic costs of nutrient acquisition and processing [11,12]. This is of crucial importance for a variety of ectotherms for which survival depends on diet composition and ambient temperature [13–16]. However, the interactive effects of temperature and diet composition on individual nutrient intake and fitness remain poorly understood, while they may provide insights to understand more global impacts at the population level [17].

The interaction between diet and temperature is particularly relevant to pollinators, such as bees, which are undergoing rapid declines driven in part by malnutrition and climate change [18,19]. Bees rely entirely on flowering plants for their nutrition, extracting mainly water and carbohydrates from nectar, as well as proteins, lipids, and many micronutrients from pollen [20]. Yet, floral nectar and pollen production are sensitive to heat and drought stress, resulting in lower-quality food [21–23]. Foraging bees must balance their nutrient collection to support their own needs, but also those of the developing brood and, in social species, the rest of the colony [24]. Bees themselves are physiologically impacted by elevating temperatures, which induce higher energy costs of foraging flights [25,26] and colony thermoregulation [27,28]. Studies using nutritional geometry, a conceptual framework to analyse the multi-dimensional food choices of animals [29,30], have shown how bees can self-select diets to achieve different nutrient balances that optimise fitness-related traits such as learning and memory [31–33], reproduction and development [34–36], immunity [37], or survival [38,39]. While this is an important first step, most of these findings currently come from experiments conducted under stable standard laboratory conditions (between 24 and 29°C), which may not reflect the thermal challenges bees face in naturally fluctuating environments.

Here, we used nutritional geometry to study how temperature influences nutrient regulation in the buff-tailed bumblebee, *Bombus terrestris*, which is particularly vulnerable to thermoregulatory challenges as it has limited homeostatic control [40] compared to other social insects. We provided micro-colonies of bees with artificial liquid diets varying in protein, carbohydrate, and lipid concentrations and ratios (Figure 1). In choice experiments, we quantified the nutrient collection of bees presented complementary diets at 30°C, a temperature considered optimal for colony development [28,41], but also at 20°C (cold) or 35°C (hot) to test whether bees can adjust nutrient regulation to a thermal stress. We then ran no-choice experiments in which bees were restricted to single diets at either 30°C or 20°C in order to assess how diet and temperature jointly affect key performance traits, and how bees navigate between these trade-offs.

**Figure 1:**
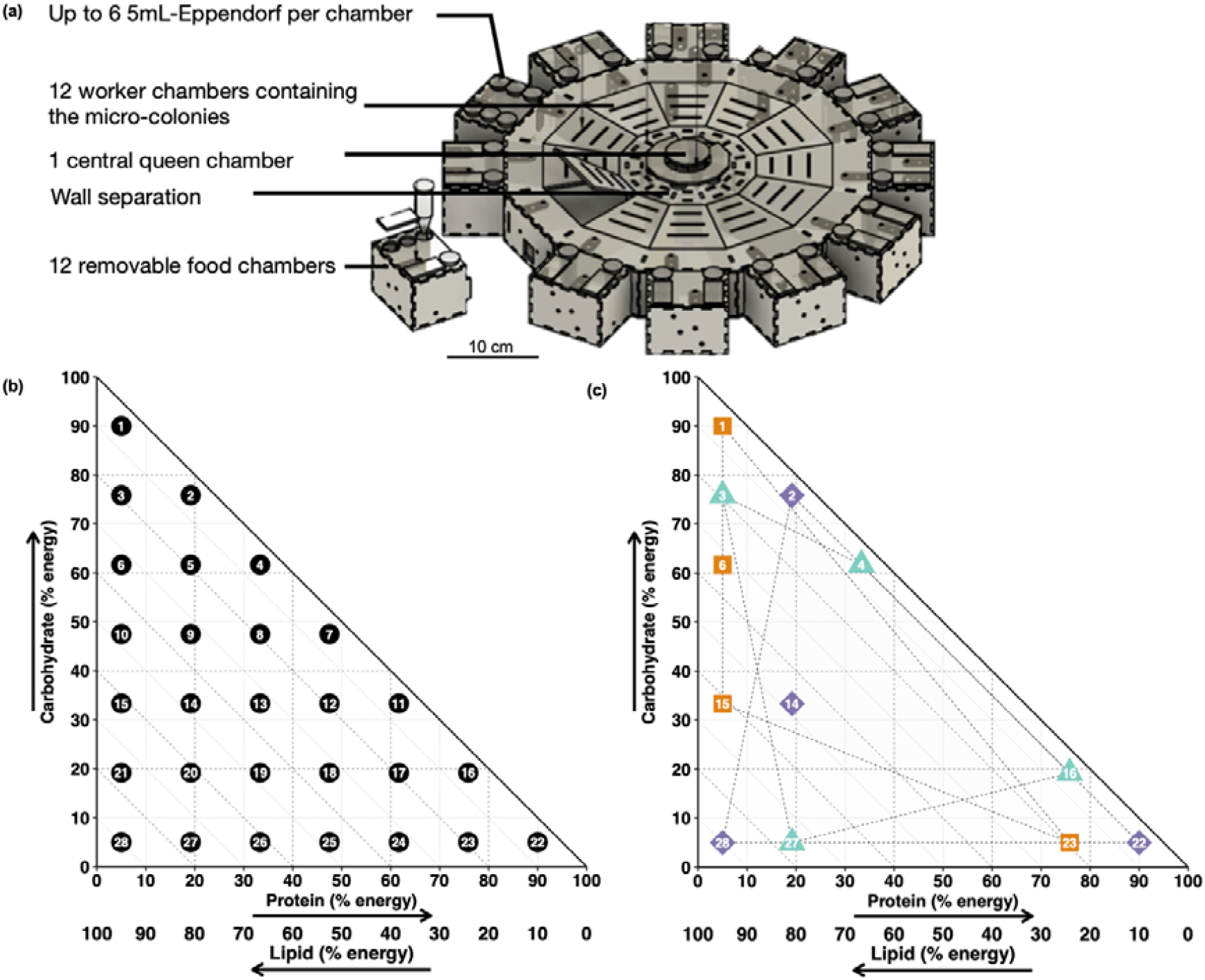
Experimental arena and 3D nutrient space representing nutritional inputs. **(a)** Dodecagonal arena used in choice and no-choice experiments; **(b-c)** 3D nutrient space in which are represented **(b)** the 28 artificial diets (numbered dots) and **(c)** the three configurations for the choice experiments (see Methods). In each configuration the bees were given a simultaneous choice between four diets (C1: orange squares, C2: purple diamonds, C3: teal triangles). See Table S1 for details about diet composition.

## Methods

### Bees

We ran the experiments between June 2020 and August 2021 with 57 commercial colonies of *B. terrestris* (Koppert BV, Berkel en Rodenrijs, The Netherlands). Each colony contained about 200 workers, brood and a queen. Upon arrival in the lab, we maintained the colonies for seven days at 30°C, with 50% relative humidity, under a 12:12h light:dark photocycle with *ad libitum* access to commercial sugar syrup (Koppert BV, Berkel en Rodenrijs, Netherlands).

### Artificial diets

We designed artificial diets varying in their contents of proteins (P), lipids (L), carbohydrates (C). We strictly used liquid diets in order to maximise the precision of food consumption measurements [42]. We used 28 diets defining a three-dimensional nutrient space (Figure 1.b). The diets had different P:C:L ratios (Table S1) but were isocaloric (0.63 cal/mg). This value was determined using the average daily food consumption of bumble bee workers obtained from previous studies (ca. 190 calories for 300 mg of diet per day: [39,43]). The caloric value of P and C was considered as 4 cal/mg and that of lipids, 9 cal/mg [44].

To make sure the nutrient content of our artificial diets resembled that of the natural diets of bees, we used the main sugars found in plant nectar [20,45]) as source of carbohydrates, namely sucrose (Interdis, Massy, France), fructose and glucose (Louis François, Paris, France). For lipids, we used sunflower oil (Auchan, Villeneuve d’Ascq, France), coconut oil (Cauvin, Saint-Gilles, France), flaxseed oil (Mon droguiste, Troyes, France), palm oil (Mon droguiste, Troyes, France) and soy lecithin (Louis François, Paris, France). For proteins, we used soy protein isolate and brown rice protein (MyProtein, Manchester, England) as well as ten essential free amino acids (valine, methionine, isoleucine, leucine, threonine, phenylalanine, lysine, histidine, arginine, tryptophan) (Fisher Scientific, Strasbourg, France). Within each diet mixture, the ratio of ingredients was calculated to match as closely as possible an average profile found in natural pollen for essential amino acids [34,37,46,47] and fatty acids [48,49]. We used xanthan and guar gums (Labo&Gato, Bordeaux, France) as emulsifiers and thickening agents to facilitate the mixing of proteins and lipids in water [50]. To prepare 1 kg of each diet, we added 1.2g of both gums in 300g of water, as well as 5g of vitamins (Vanderzant vitamins mix, Sigma-Aldrich, Darmstadt, Germany). We then added all other ingredients with the correct amount of remaining water. All diets had nutrient concentrations (proteins: 1.5-48.4%; lipids: 1.2-24.6%; carbohydrates: 10-70%) falling within the range of natural pollens and nectars [20,51–53]. We mixed the diets using a blender and stored them at -20°C until usage. Preliminary observations showed that bumblebees successfully collected the different diets and stored them in empty brood cells (i.e. honey pots), corroborating the suitability of our diets for experiments of nutritional regulation (Figure S1). In what follows, we used the terminology ‘diet’ to refer to these artificial food sources characterised by specific nutrient ratios and ‘food’ to refer to the total quantity of nutrients collected by the bees.

### Behavioural assay

We ran experiments on micro-colonies of bee adult workers. Each of the 57 mother colonies was divided in 6 to 12 micro-colonies, containing 20 workers and ca. 77.98g (± 10.49 CI95) of empty brood cells, acting as food reserves, from the mother colony. The bees were placed in a homemade designed dodecagonal arena (Figure 1.a) enabling for the physical separation between groups of workers and their queen while maintaining chemical communication involved in social cohesion [54] (see details in Figure S2). The colony queen was placed in the central chamber containing brood cells and fed commercial sugar syrup (Koppert BV, Berkel en Rodenrijs, Netherlands). The micro-colonies built with workers from the same mother colony were placed in peripheral chambers of the arena and fed artificial diets.

Micro-colonies were maintained at 50% relative humidity, under a 12:12h light:dark photocycle, for 14 days. During this period, we measured the following performance traits: diet collection, total food collection, workers survival, adult relative dry weight (dry weight divided by thorax width [55]) (proxy for energy reserves [56]), and egg-laying (proxy for fecundity [57]). We provided artificial diets every day in drilled 5ml Eppendorf vials used as gravity feeders (Figure S1), containing 4.36g (± 0.01 CI95). We weighed each vial before introduction and after removal from food chambers, using a precision balance of 1mg (ME103T, Mettler Toledo, Greifensee, Suisse). In parallel, we placed vials in similar empty boxes to evaluate evaporation. Time between each diet renewal was measured. Dead bees were counted, removed, and frozen for later analyses every day. The brood cells placed in each micro-colony were dissected at the end of the experiment to count the number of eggs and larvae. Pupae were not included because they would have been produced by the mother colony (egg to pupa development takes between 13 and 26 days depending on the sex and nest condition [58]) and undetected when forming the micro-colony.

### Choice experiments

We assessed the nutrient intake target - the balanced ratio of carbohydrates, proteins, and lipids [59,60] - of bees by giving them a simultaneous choice of four diets at three temperature regimes: 20°C (cold for the bees), 30°C (reference temperature for optimal colony development) and 35°C (hot for the bees). Room temperature was measured every five minutes using a data logger (Elitech, London, United Kingdom) and remained in the range of 0.01°C of the desired temperature during the entire experimental duration. At each temperature, micro-colonies were given a choice of four complementary diets, out of three possible combinations (C1: 1, 6, 15, 23; C2: 2, 14, 22, 28; C3: 3, 4, 16, 27) (see diet combinations in nutrient space in Figure 1.c). Using multiple diet combinations enabled us to discriminate between selective diet collection and random diet collection (which would systematically correspond to the geometrical barycentre of each combination in the 3D nutrient space). We tested 8 micro-colonies per choice configuration at 20°C and 30°C, and 16 micro-colonies per choice configuration at 35°C. We made more replicates at 35°C to compensate for the higher mortality rates (see Results).

### No-choice experiments

We assessed the consequences of different nutrient collections on fitness and investigated the nutritional rules of compromise - the patterns of feeding in imbalanced diets [6]-used by bees to mitigate the effects of feeding away from their intake target. These experiments were run only at 20°C and 30°C. When trying to run our experiment at 35°C, 50% of the bees were dead as soon as day 3, which greatly limited sample size to provide meaningful data. For the no-choice experiments at 20°C and 30°C, bees were provided two gravity feeders of 6.25g (± 0.05 CI95) of the same diet. All 28 diets were used at 30°C, whereas only the 10 diets promoting higher survival at 30°C were used at 20°C, here also to provide enough data. Some micro-colonies were only fed on water at 20°C or 30°C to serve as controls for survival, weight and egg-laying comparisons. No-choice experiments were replicated eight times at 20°C and seven times at 30°C.

### Measure of food collection

We measured the average amounts of total food and of each specific macronutrient collected per bee per day as follows.

The initial food weight (IF) was calculated with:

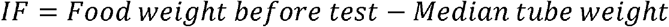

where food weight before the test is the weight of the full tube, and the median tube weight (tare) was calculated from 100 empty tubes.

The remaining food weight (RF) was calculated with:

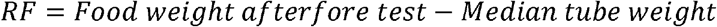

where food weight after test is the weight of the full tube after the experimental period. To account for evaporation, RF was corrected by subtracting the mean evaporative loss from identical tubes placed in empty boxes over the same period.

We then calculated the daily collected food per bee (DF) as:

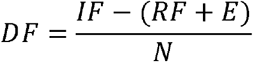

where E is the mean evaporative loss per tube and N is the number of bees.

The cumulative collected food per bee (CF) was next calculated as:

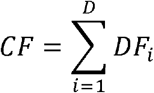

where DF_i_ is the daily collected food on day i, summed over all experimental days D.

Finally, we calculated the cumulative amount (CN) of each nutrient collected per bee as follows:

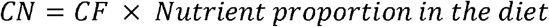

where the nutrient proportion depended on the experimental diet (Table S1), and was expressed either as caloric contribution (energy consumed) or as mass fraction (mg of nutrient).

### Body size and weight

As body size affects foraging efficiency and division of labour in bumblebees [61], we controlled if it had an influence on nutrient regulation. After the behavioural tests, all bees were frozen to measure their thorax width (intertegular span) as a proxy for overall body size, using a binocular microscope (Nikon Europe B.V., Amsterdam, Netherlands). Bees were then dried at 80°C during 4h in an oven (WTB Binder, Tuttlingen, Germany) before being weighed using a precision balance of 1mg (ME103T, Mettler Toledo, Greifensee, Switzerland).

### Statistical analysis

The raw data used in this study are available in the online repository Zenodo (https://doi.org/10.5281/zenodo.18339387). Data are presented in the text as median with the limits of 95% confidence intervals into brackets. All statistical analyses were performed in R Studio 2024.04.24 [62]. We normalised and transformed the variables (e.g. relative dry weight (in mg) for a median bee (of 4.5mm thorax width), the brood size (number eggs and larvae) found at the end of experiments) using the formula:

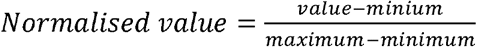

We then removed zeros and ones using:

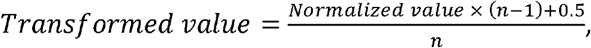

where n is the number of values [63].

We used general linear mixed models (GLMM) from package Template Model Builder (version 1.1.9) (Brooks et al., 2017), day and colony identity nested in replicate as random factors. For the choice experiments, we set groups of choice configuration, temperature, and their interaction as fixed effects, and for the no-choice experiments, only the diet was set as fixed effect. GLMM were followed by an Anova.glmmTMB function from that package to assess the overall significance of the fixed effects, and we then ran post-hoc comparisons using package emmeans (version 1.10.2) [64]. We performed survival analyses using Cox Proportional-Hazards Models (version 2.2-22) [65] on groups in choice and no-choice experiments, with the colony identity as random factor, followed by pairwise comparisons using Hazard Ratios. We performed linear programming using lpSolve package (version 5.6.23) [66] to find the minimum amount of food mixture satisfying the given constraints on total food consumption, quantity of nutrients and water.

## Results

### Bees collected more food with increasing temperature

Since the artificial diets were the only source of water to the bees, we first tested whether workers adjusted their water collection by analysing nutrient to water (N:W) regulation in the choice experiments. N:W collection by individual bees was influenced by the interaction between choice configuration and temperature (ANOVA: Configuration*Temperature: Chi=49.147, df=4, p<0.001). Bees thus collected significantly more water with increasing temperatures (Figure 2.a), with a seven-fold augmentation between the three temperature conditions, from a median of 55.03 mg/bee/day [95CI: 51.57; 69.11] at 20°C to 140.79 mg/bee/day [129.97; 173.47] at 30°C, and 399.25 mg/bee/day [364.03; 469.91] at 35°C (Figure 2.a).

**Figure 2:**
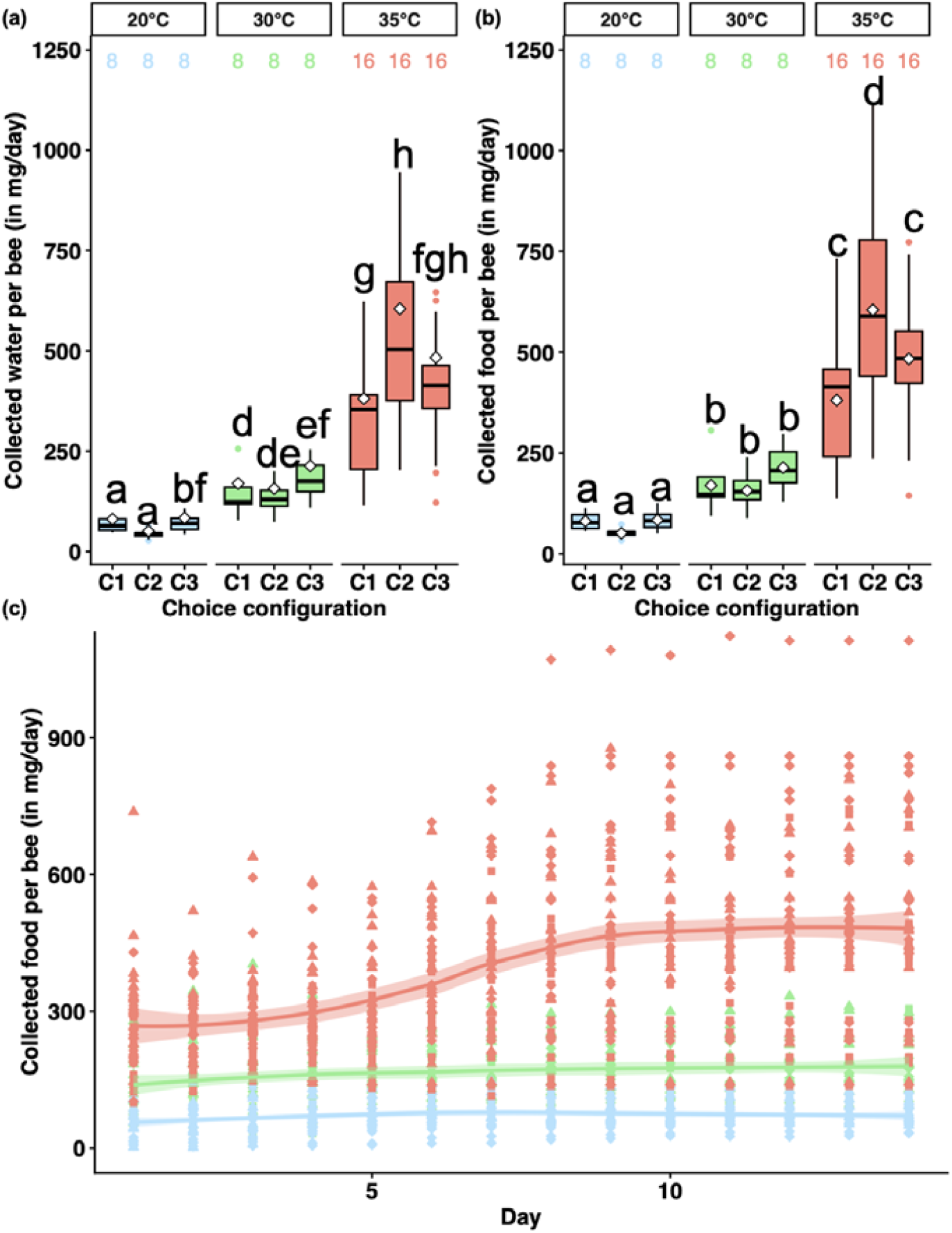
Food and water collection for each temperature and choice configuration. **(a)** Boxplot of total water and **(b)** food collected per bee (in mg/day) in the choice experiments, for each temperature and each choice configuration. The boxplots display the mean (white diamond) and median (central line), the edges of the box are the 25th and 75th percentiles. The whiskers extend to the most extreme data points without considering outliers, which are represented by dots. Letters (a-f) show significant differences from GLMM. **(c)** Food collection over time in the choice experiments. Fitted lines for temperature are displayed with 95% confidence intervals. Dot shapes (square, diamond or triangle) correspond to the diet (C1-C3).

In order to focus on nutrient collection, we added N:W as a random factor in all the subsequent analyses. We found a significant interaction between choice configuration and temperature on food collection (ANOVA: Configuration*Temperature: Chi=19.903, df=4, p<0.001). Overall, bees collected more food (total nutrients) as temperature increased, from a median of 65.05 mg/bee/day [95CI: 52.41; 95.8] at 20°C to 166.43 mg/bee/day [136.98; 238.9] at 30°C, and 467.29 mg/bee/day [321.67; 647.42] at 35°C (Figure 2.b). We observed the same trend when considering only the combined collection of proteins, carbohydrates and lipids (ANOVA: Configuration*Temperature: Chi=19.930, df=4, p<0.001). This impact of temperature was different across conditions as indicated by the day-to-day pattern of food collection by bees (Figure 2.c). At 20°C, there was no interaction between temperature and day, the collection of food increased only slightly over time (from 61.89 mg/bee [44.22; 74.29] on the first day, to 65.05 mg/bee [61.41; 82.05] on the last day), and followed a pattern comparable to that observed in the reference temperature of 30°C (GLMM: Temperature x Day: 0.001±0.009, p=0.903). At 30°C, the daily food collection increased steadily with time (from 120.11 mg/bee [99.50; 166.2] on the first day, to 166.43 mg/bee [154.56; 205.65] on the last day) (GLMM: Temperature x Day: 0.031±0.007; p<0.001). However, at 35°C, the food collection started at a higher level than at 20 or 30°C (260.65 mg/bee [239.33; 301.55] on the first day) and increased with time (467.29 mg/bee [427.75; 551.52] on the last day) (GLMM: Temperature x Day: 0.029±0.007, p<0.001).

### Bees increased the P:C ratio in collected food with increasing temperatures

We next investigated the impact of temperature on the mean nutrient collection by the bees in the choice experiments (Table 1). Temperature significantly influenced the proportions of collected carbohydrates (ANOVA: Chi=7.007, df=2, p=0.030; Figure 3.a) and proteins (ANOVA: Chi=40.387, df=2, p<0.001; Figure 3.b). Specifically, the proportion of carbohydrates decreased between 20°C and 35°C (GLMM: p=0.028), while that of proteins increased (GLMM: p<0.001 for all pairwise comparisons, except between 30°C and 35°C: p=0.063). By contrast, the mean proportion of lipids collected by the bees remained unaffected by temperature (Chi=0, df=2, p=1; Figure 3.c). This suggests that bees prioritised lipid regulation over that of the two other nutrients (Figures 3.d and 3.e). For all the diet choice configurations, the random proportions of intake for all the nutrients did not fall within the 95CI of the real nutrient intake experimentally measured (Table 1), indicating that bees made active (non-random) nutrient balancing by selectively collecting the available diets.

**Table 1:**
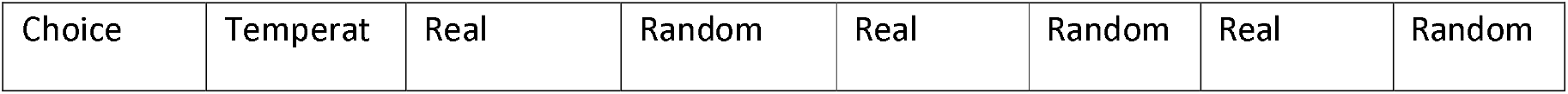
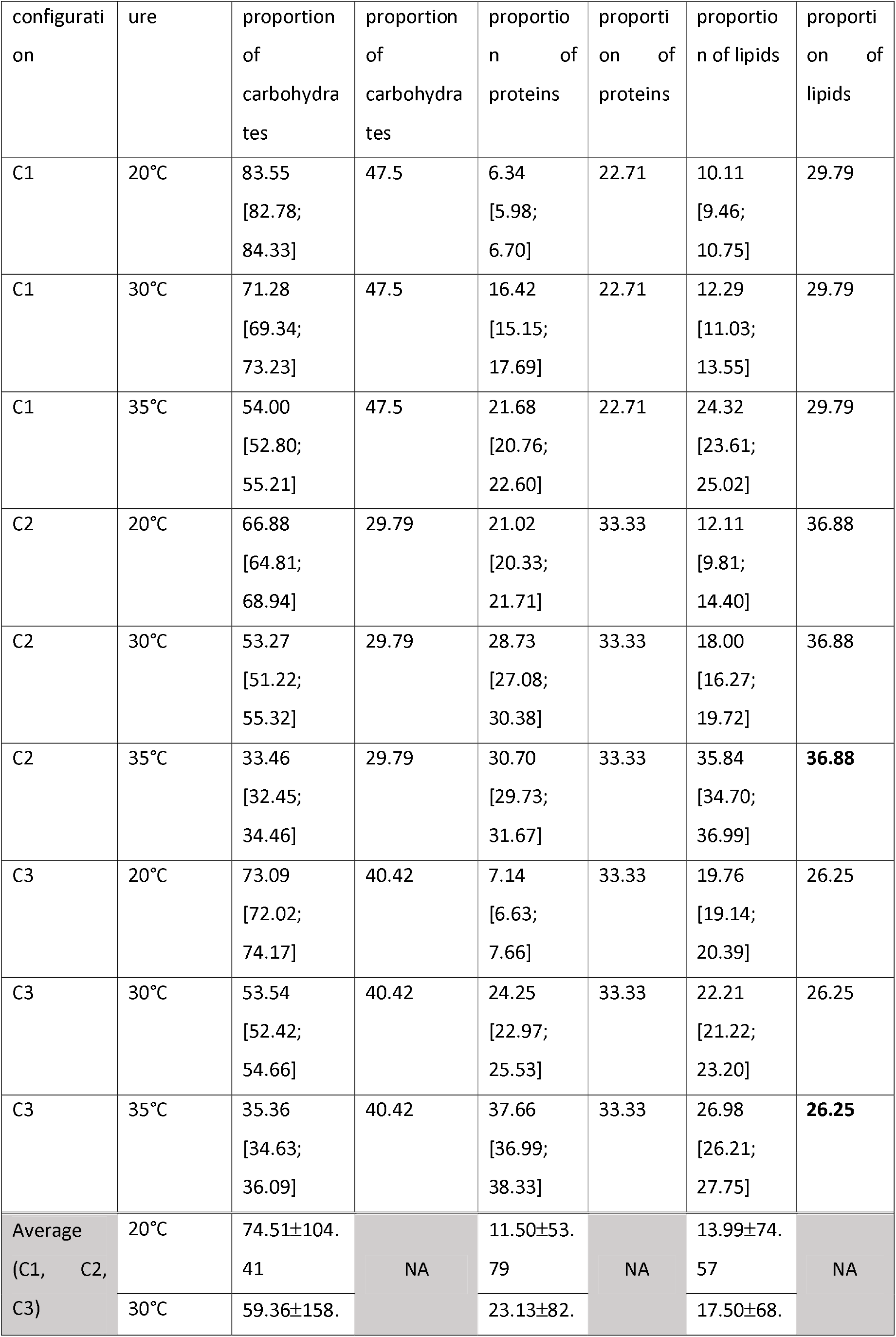
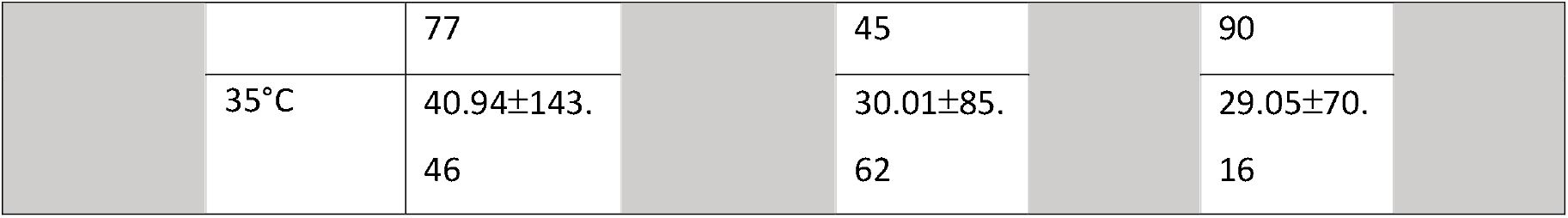
Mean proportion of nutrients collected. Upper panel: mean [95CI] proportions of nutrients collected per diet and temperature and comparison with the random intake. The random collection value is displayed in bold when included within the 95CI of the real data. Lower panel: mean (±variance) proportions of collected nutrients at each temperature (see Figure 3). NA: not applicable.

|  |  |  |  |  |  |  |  |
| --- | --- | --- | --- | --- | --- | --- | --- |
| Choice | Temperat | Real | Random | Real | Random | Real | Random |

| configuration | ure | proportion<br>of<br>carbohydrates | proportion<br>of<br>carbohydrates | proportion<br>of<br>proteins | proportion<br>of<br>proteins | proportion<br>of lipids | proportion<br>of<br>lipids |
| --- | --- | --- | --- | --- | --- | --- | --- |
| C1 | 20°C | 83.55<br>[82.78;<br>84.33] | 47.5 | 6.34<br>[5.98;<br>6.70] | 22.71 | 10.11<br>[9.46;<br>10.75] | 29.79 |
| C1 | 30°C | 71.28<br>[69.34;<br>73.23] | 47.5 | 16.42<br>[15.15;<br>17.69] | 22.71 | 12.29<br>[11.03;<br>13.55] | 29.79 |
| C1 | 35°C | 54.00<br>[52.80;<br>55.21] | 47.5 | 21.68<br>[20.76;<br>22.60] | 22.71 | 24.32<br>[23.61;<br>25.02] | 29.79 |
| C2 | 20°C | 66.88<br>[64.81;<br>68.94] | 29.79 | 21.02<br>[20.33;<br>21.71] | 33.33 | 12.11<br>[9.81;<br>14.40] | 36.88 |
| C2 | 30°C | 53.27<br>[51.22;<br>55.32] | 29.79 | 28.73<br>[27.08;<br>30.38] | 33.33 | 18.00<br>[16.27;<br>19.72] | 36.88 |
| C2 | 35°C | 33.46<br>[32.45;<br>34.46] | 29.79 | 30.70<br>[29.73;<br>31.67] | 33.33 | 35.84<br>[34.70;<br>36.99] | **36.88** |
| C3 | 20°C | 73.09<br>[72.02;<br>74.17] | 40.42 | 7.14<br>[6.63;<br>7.66] | 33.33 | 19.76<br>[19.14;<br>20.39] | 26.25 |
| C3 | 30°C | 53.54<br>[52.42;<br>54.66] | 40.42 | 24.25<br>[22.97;<br>25.53] | 33.33 | 22.21<br>[21.22;<br>23.20] | 26.25 |
| C3 | 35°C | 35.36<br>[34.63;<br>36.09] | 40.42 | 37.66<br>[36.99;<br>38.33] | 33.33 | 26.98<br>[26.21;<br>27.75] | **26.25** |
| Average<br>(C1, C2,<br>C3) | 20°C | 74.51±104.<br>41 | NA | 11.50±53.<br>79 | NA | 13.99±74.<br>57 | NA |
|  | 30°C | 59.36±158. |  | 23.13±82. |  | 17.50±68. |  |

|  |  |  |  |  |  |  |
| --- | --- | --- | --- | --- | --- | --- |
|  |  | 77 |  | 45 |  | 90 |
|  | 35°C | 40.94±143. |  | 30.01±85. |  | 29.05±70. |
|  |  | 46 |  | 62 |  | 16 |

**Figure 3:**
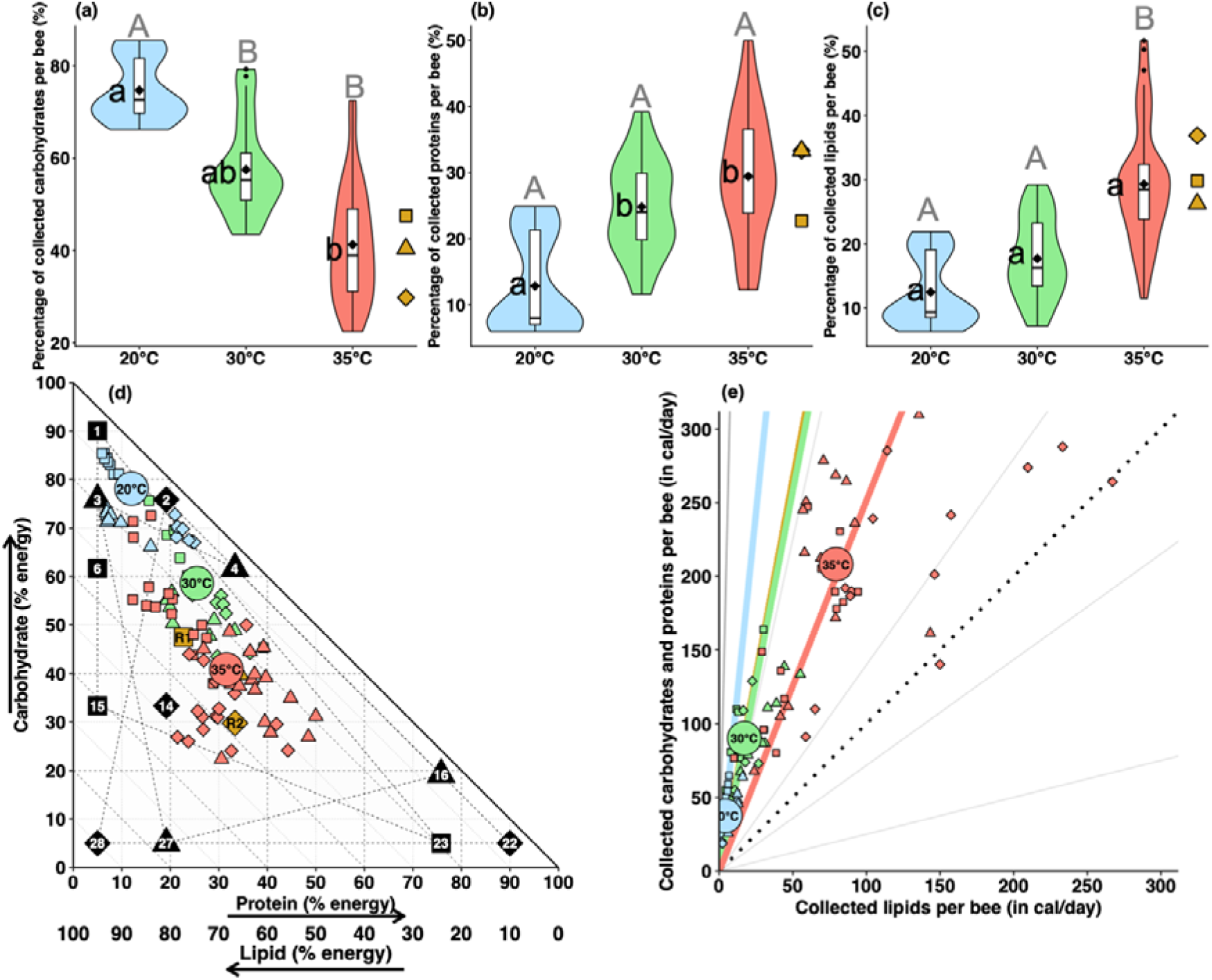
Proportion of nutrients collected in the choice experiments. **(a-c)** Violin and boxplots of total nutrients collected per bee (in %) at the end of the choice experiment, for each temperature. The boxplots display the mean (black diamond) and median (central line), the edges of the box are the 25th and 75th percentiles. The whiskers extend to the most extreme data points without considering outliers, which are represented by dots. The violin plots wrapping outside the boxplots show the data distribution. The proportions expected from random choices are displayed in yellow, with shapes (square, diamond or triangle) corresponding to the diet. Lowercase letters next to boxes (a-c) show significant differences between means from GLMM. Grey uppercase letters above boxes (A-B) show significant differences between variances from Levene’s test. **(d)** 3D nutrient space, where the dotted polygons are the three choice configurations with their respective four diets (numbered squares, diamonds or triangles) and corresponding random points (yellow square, diamond or triangle). In all panels, the coloured squares, diamonds or triangles indicate the proportions of nutrients collected by each individual at the end of the experiment, while coloured circles indicate the median for each group (20°C, 30°C, 35°C). **(e)** Nutrient geometry plot representing amounts of lipids vs. carbohydrates + proteins collected by the bees. Grey lines: ratio in each diet; yellow line: ratio expected for a random choice; coloured lines: median ratio obtained for each group; dotted line: 1:1 ratio. Coloured squares, diamonds or triangles indicate the proportions collected by each individual at the end of the experiment. Coloured circles indicate the median for each group (20°C, 30°C, 35°C).

When looking at the impact of temperature on the variability of nutrient collection by the bees, variance for protein collection remained stable across temperature regimes (Figure 3.b). Variance for carbohydrates collection increased between 20°C and the two higher temperatures (Bartlett’s test: p<0.05 for pairwise comparisons; Figure 3.a), but remained stable between 30°C and 35°C (Bartlett’s test: p=0.255). Variance for lipid collection was similar between 20°C and 30°C (Bartlett’s test: p=0.394) and increased at 35°C (Bartlett’s test: p=0.001 between 20°C and 35°C; p=0.044 between 30 and 35°C). Thus overall, higher temperatures increased variability in carbohydrate and lipid collection. The reduced accuracy in nutrient balancing by the bees exposed to 35°C was associated to significantly reduced fitness traits such as survival and body weight, as compared to at 20°C and 30°C (Figure S3, Text S1).

### Bees prioritised survival over body condition at 20°C

We measured workers’ survival, relative weight and egg-laying in the no-choice experiments, to identify whether any of these performance traits were prioritised over others, at the reference temperature of 30°C or under a cold stress at 20°C. We did not analyse the data for 35°C as 50% of the bees died after only three days.

When constrained to a single diet, the survival probability of bees was influenced by the diet at 30°C (ANOVA: Chi=143.54, df=28, p<0.001) and at 20°C (ANOVA: Chi=26.363, df=10, p=0.003). At both temperatures, the best survival performances were found for carbohydrate-rich diets (mostly diets 2 and 3) while the worst ones were found for protein-rich diets (Figure 4.a-b). The relative dry weight (in mg) for a median-sized bee (4.5mm thorax width) was also influenced by the diet, at both 30°C (ANOVA: Chi=58.354, df=28, p<0.001) and 20°C (ANOVA: Chi=35.608, df=10, p<0.001). For both temperatures, bees collecting carbohydrate-rich diets (mostly diets 1, 2, 4 and 5) were larger than those collecting carbohydrate-poor diets (Figure 4.c-d). As the brood count data did not meet the quality standards required for reliable analysis (64% of the micro-colonies had no egg laid during the experiment), they were excluded from the analysis.

**Figure 4:**
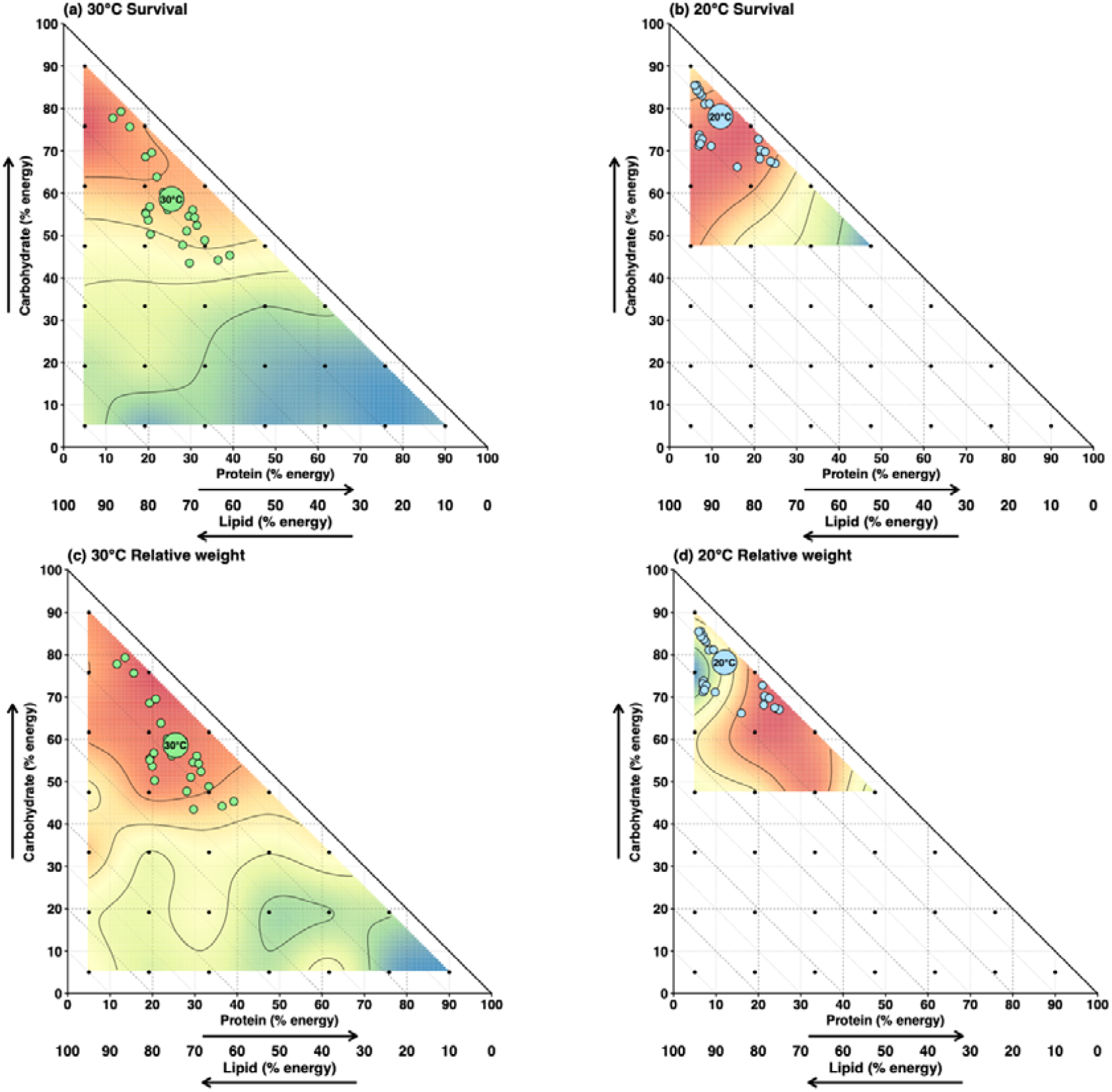
Diet collection in choice experiments mapped on the performance traits measured in no-choice experiments. Heatmaps on 3D nutrient space representing: **(a, b)** adult survival and **(c, d)** the relative dry weight (in mg) for a median-sized bee (4.5mm thorax width) at 30°C (left panels) and 20°C (right panels). Warmer colours indicate better performances. Small coloured circles represent nutrient collection from each micro-colony in choice experiments, while large coloured circles represent their medians. All 28 diets were used at 30°C, whereas only the 10 diets promoting higher survival at 30°C were used at 20°C, hence the cropped landscapes at 20°C.

Based on the above observations, we next created performance landscapes (Figure 4). The shape and position of performance peaks for survival (Figure 4.a-b) and relative weight (Figure 4.c-d) differed between temperatures. Plotting the nutrient collection data on these performance landscapes informed about how bees navigate potential trade-offs between the different fitness traits. At 30°C, bees collected nutrients in proportions that optimised both their survival (Figure 4.a) and body weight (Figure 4.c). However, at 20°C, diet collection overlapped with the peak of survival (Figure 4.b) but not with that of body weight (Figure 4.d). Together, these analyses suggest that bees navigate fitness trade-offs differently depending on temperature, prioritising survival over weight in colder, suboptimal conditions for colony development.

## Discussion

Using a 3D nutritional geometry design, we showed how temperature profoundly influences the quantity of food collected by bumblebees, their strategies of nutrient regulation, and ultimately their fitness. At 30°C, bees successfully balanced carbohydrate, protein, and lipid collection, at levels beneficial for body weight and survival. At 20°C, bees reduced their overall nutrient collection, selecting proportionally more carbohydrates while keeping lipid levels constant, thereby prioritising immediate survival while losing weight. At 35°C, however, nutrient balancing was disrupted and survival dropped dramatically.

### High temperature alters intake demand and hydration regulation

Bees markedly increased both food and water collection at elevated temperature (35°C), which is consistent with elevated metabolic demands and increased risk of dehydration at higher temperatures [67,68]. While bumblebees have anecdotally been observed collecting water for thermoregulation under warm temperature [69], it is believed that most of their water needs is extracted from nectar. In our experiments, as water was only available through the liquid diets, the strong increase in water intake indicates active regulation of hydration rather than a passive consequence of increased feeding. This result aligns with nutritional geometry theory, which predicts that organisms should adjust collection of key food components to meet changing physiological requirements when environmental constraints shift [6].

Notably, food collection at an elevated temperature of 35°C followed a distinct temporal pattern, characterised by high initial collection rates followed by increased collection and variability. This suggests that although bumblebees initially compensated for thermal stress by increasing food collection, sustained regulation became difficult under extreme heat. Such early over-collection is consistent with the idea that physiological limits constrain nutrient regulation under stressful conditions. Consistent with this interpretation, recent studies show that bumblebee workers fail to maintain effective thermoregulation during short-term heatwave exposure, leading to impaired physiological and sensory performances, even though heatwave exposure during development does not affect adult body size [40].

### Temperature shifts nutrient intake targets without abolishing regulation

Exposure to different ambient temperatures significantly altered bees’ nutrient balancing. When exposed to high temperatures, bees shifted their intake nutrient collection target toward higher protein and lower carbohydrate proportions, while lipid collection remained remarkably stable. This indicates temperature modifies the position of bees’ collection target in the nutrient space rather than relaxing their nutrient regulation altogether. A stable proportion of lipid intake across temperatures suggests strong prioritisation of lipid regulation over other macronutrients, consistent with other recent evidence [49]. Importantly, this prioritisation persisted even under thermal stress, when regulatory precision for carbohydrates—and to a lesser extent proteins—declined substantially. Bees thus tolerated increased variance in carbohydrate intake and reduced fitness at high temperature before deviating from lipid targets, indicating that lipid balance is subject to tighter physiological constraints than the other macronutrients. At all tested temperatures, bees regulated their lipid intake in a non-random manner, suggesting an inherent limit to the lipid appetite. This pattern fits well with recent work by Ruedenauer et al. [70], who demonstrated that bumblebees prioritise lipids when evaluating pollen taste, highlighting lipid regulation as a key determinant of foraging behaviour [71,72].

This strong regulation of lipid intake may reflect its multifunctional role in bee physiology [35,73], related to membrane stability, energy storage, or signaling functions that are critical across thermal environments. Recent work has also highlighted sterol availability as a critical and underappreciated constraint on bee nutrition [74], with consequences for development, survival, and resilience to environmental stress. From this perspective, the tight regulation of lipid collection observed in our study may represent an indirect but essential mechanism for securing adequate sterol supply, especially under elevated temperatures where membrane stability and cellular maintenance demands are heightened. In contrast, the opposing shifts in protein and carbohydrate collection imply a temperature-dependent rebalancing of metabolic demands, possibly reflecting increased protein requirements for cellular maintenance and repair at higher temperatures.

Importantly, nutrient intake differed from random expectations across all diet configurations, confirming that bees in our experiments actively regulated nutrient collection even under thermal stress. Thus, temperature does not eliminate nutrient regulation, but triggers changes in the rules governing nutrient collection decisions by bumblebees. Note however that increased temperature reduced the precision with which nutrients were balanced. In particular, variance in nutrient collection increased substantially at higher temperatures at 35°C. From a nutritional geometry perspective, increased variance reflects a widening of intake trajectories around the target, indicative of reduced regulatory accuracy. Such imprecision associated with reduced survival and body weight at 35°C suggests nutrient regulation was close to complete disruption at such high temperature.

### Temperature determines how nutritional trade-offs are navigated

Mapping the results of the choice experiments on performance landscapes shows that temperature alters how bumblebees resolve trade-offs between fitness components. At 30°C, individuals collected nutrients in proportions that simultaneously optimised survival and body weight, indicating alignment of nutritional optima under mean conditions. In contrast, at 20°C, bees collected carbohydrate-rich diets that maximised survival (notably thermogenesis [20]) at the expense of body mass, suggesting a prioritisation of immediate survival under cold stress. This strategy parallels the overwintering behaviour of bumblebee queens that considerably reduce their activity and metabolic rates to conserve energy, while consuming internal reserves [75–78]. This temperature-dependent shift in prioritisation is a central prediction of nutritional geometry: when multiple performance surfaces do not align, animals should target the trait most critical for immediate fitness. Our results provide empirical support for this, showing how bumblebees flexibly adjust their nutritional strategy depending on environmental context.

### Implications for pollinator resilience under climate change

Our findings in tightly controlled laboratory conditions have potentially important implications for understanding pollinator responses to climate change. Rising temperatures are likely to increase energetic and hydration demands of bumblebees while simultaneously reducing the precision of nutrient regulation [79,80]. If floral resources do not match temperature-dependent nutritional requirements, bumblebees may be forced to navigate into suboptimal regions of the nutrient space, exacerbating fitness costs. Moreover, floral resources could also be impacted by elevated temperature, with hot and dry conditions reducing nectar volume in flowers [21–23]. The observed breakdown of regulatory precision of bumblebees at high temperature suggests that nutritional plasticity has limits, beyond which individuals may be unable to compensate for environmental stress. Social insects can buffer some environmental variation through nest thermoregulation and behavioural adjustments [81], yet the increasing frequency, amplitude and duration of heatwaves [82,83] may overwhelm such resilience. Under increasingly frequent thermal extremes, such constraints could contribute to reduced survival, impaired condition, and ultimately population-level consequences.

## Conclusions

Using 3D nutritional geometry experiments, we showed that temperature stress influences bumblebee’s nutrient collection targets, regulatory precision and expression of fitness traits. Temperature stress scale food collection and fundamentally alters how bumblebees navigate their nutrient space and trade-offs between fitness traits. Given their role as essential ecosystem service providers through pollination, our study uncovers the underappreciated effects of temperature stress on nutritional needs and choices that can determine how bees respond to changing climates.

## Acknowledgements

We thank Julie Peuzé for her help with data collection during a Master internship. We are also grateful to Camille Buhl for fruitful discussions on an earlier previous draft. This work was funded by a PhD grant from the Association Nationale Recherche Technologie (ANRT) to Koppert and the CNRS. While writing, CM and ML were funded by an ERC-Consolidator grant to (BEE-MOVE GA101002644 to ML).

## Authors contribution

CM: Formal analysis, Validation, Visualization, Roles/Writing - original draft, Writing - review & editing; SK: Data curation, Formal analysis, Validation, Roles/Writing - original draft, Investigation, Methodology, Software; JG: Supervision, Writing - review & editing; JMD: Supervision, Writing - review & editing; JM: Formal analysis, Visualization, Writing - review & editing; ML: Project administration, Resources, Supervision, Funding acquisition, Methodology.

## Competing interests

The authors declare no competing interests.

## Notes

https://doi.org/10.5281/zenodo.18339387

### Competing Interest Statement

The authors have declared no competing interest.

