## Supplementary Information for "Nutritional responses of bumblebees to thermal stress"

This file contains:

Table S1: Recipes of the 28 artificial diets.

Figure S1: Collection of artificial diet by bumblebees.

Figure S2: Technical schematics of the dodecagonal arena used in the choice and no-choice experiments.

Figure S3: Fitness performances for the three temperature regimes in the choice experiment.

Text S1: Fitness performances in the choice experiments.

**Table S1: Recipes of the 28 artificial diets**. % of each nutrient (cal): Percentages of carbohydrates (C) (in red), lipids (L) (in green) and proteins (P) (in blue) in each diet in calories. Nutrient in diet (g/kg): Amounts of carbohydrates, lipids, proteins, total nutrients, and water in grams per kilograms. Ratio N:W: ratios of all nutrients (N) to water (W). Ratio nutrient1 (nutrient2+nutrient3): ratios of each macronutrient to the two others, in calories. Macronutrient mixtures compositions are described in the Methods. Diets displayed with a grey background were used in the choice experiment.

| Diets | % C | % L | % P | C in diet  (g/kg) | L in diet  (g/kg) | P in diet  (g/kg) | **Total Nutrient (g/kg)** | **Water (g/kg)** | **Ratio N:W** | Ratio C:(L+P) | Ratio L:(C+P) | Ratio P:(C+L) | C Mix (g/kg) | L Mix (g/kg) | P Mix (g/kg) |
| --- | --- | --- | --- | --- | --- | --- | --- | --- | --- | --- | --- | --- | --- | --- | --- |
| **Diet 1** | **90.00** | **5.00** | **5.00** | **142.50** | **3.51** | **7.91** | **153.93** | **835.60** | **5.43:1** | **09:01** | **01:19** | **5.43:1** | **144.15** | **3.55** | **9.27** |
| **Diet 2** | **75.83** | **5.00** | **19.16** | **120.06** | **3.51** | **30.34** | **153.93** | **832.02** | **5.41:1** | **3.14:1** | **01:19** | **1:4.22** | **121.46** | **3.55** | **35.55** |
| **Diet 3** | **75.83** | **19.16** | **5.00** | **120.06** | **13.48** | **7.91** | **141.47** | **848.21** | **06:01** | **3.14:1** | **1:4.22** | **01:19** | **121.46** | **13.64** | **9.27** |
| **Diet 4** | **61.66** | **5.00** | **33.33** | **97.63** | **3.51** | **52.77** | **153.93** | **828.24** | **5.38:1** | **1.61:1** | **01:19** | **01:02** | **98.77** | **3.55** | **61.83** |
| Diet 5 | 61.66 | 19.16 | 19.16 | 97.63 | 13.48 | 30.34 | 141.47 | 844.62 | 5.97:1 | 1.61:1 | 1:4.22 | 1:4.22 | 98.77 | 13.64 | 35.55 |
| **Diet 6** | **61.66** | **33.33** | **5.00** | **97.63** | **23.45** | **7.91** | **129.01** | **860.81** | **5.67:1** | **1.61:1** | **01:02** | 01:19 | **98.77** | **23.73** | **9.27** |
| Diet 7 | 47.50 | 5.00 | 47.50 | 75.20 | 3.51 | 75.20 | 153.93 | 824.84 | 6.36:1 | 1:1.11 | 01:19 | 1:1.11 | 76.08 | 3.55 | 88.1 |
| Diet 8 | 47.50 | 19.16 | 33.33 | 75.20 | 13.48 | 52.77 | 141.47 | 841.04 | 5.94:1 | 1:1.11 | 1:4.22 | 01:02 | 76.08 | 13.64 | 61.83 |
| Diet 9 | 47.50 | 33.33 | 19.16 | 75.20 | 23.45 | 30.34 | 129.01 | 857.23 | 6.64:1 | 1:1.11 | 01:02 | 1:4.22 | 76.08 | 23.73 | 35.55 |
| Diet 10 | 47.50 | 47.50 | 5.00 | 75.20 | 33.42 | 7.91 | 116.55 | 873.42 | 7.49:1 | 1:1.11 | 1:1.11 | 01:19 | 76.08 | 33.81 | 9.27 |
| Diet 11 | 33.33 | 55 | 61.66 | 52.77 | 3.51 | 97.63 | 153.93 | 821.26 | 5.34:1 | 01:02 | 01:19 | 1.61:1 | 53.39 | 3.55 | 114.38 |
| Diet 12 | 33.33 | 19.16 | 47.50 | 52.77 | 13.48 | 75.20 | 141.47 | 837.65 | 5.92:1 | 01:02 | 1:4.22 | 1:1.11 | 53.39 | 13.64 | 88.1 |
| Diet 13 | 33.33 | 33.33 | 33.33 | 52.77 | 23.45 | 52.77 | 129.01 | 853.64 | 6.62:1 | 01:02 | 01:02 | 01:02 | 53.39 | 23.73 | 61.83 |
| **Diet 14** | **33.33** | **47.50** | **19.16** | **52.77** | **33.42** | **30.34** | **116.55** | **869.83** | **7.46:1** | **01:02** | **1:1.11** | **1:4.22** | **53.39** | **33.81** | **35.55** |
| **Diet 15** | **33.33** | **61.66** | **5.00** | **52.77** | **43.39** | **7.91** | **104.08** | **886.03** | **8.51:1** | **01:02** | **1.61:1** | **01:19** | **53.39** | **43.9** | **9.27** |
| **Diet 16** | **19.16** | **5.00** | **75.83** | **30.34** | **3.51** | **120.06** | **153.93** | **817.67** | **5.31:1** | **1:4.22** | **01:19** | **3.14:1** | **30.70** | **3.55** | **140.66** |
| Diet 17 | 19.16 | 19.16 | 61.66 | 30.34 | 13.48 | 97.63 | 141.47 | 833.86 | 5.89:1 | 1:4.22 | 1:4.22 | 1.61:1 | 30.70 | 13.64 | 114.38 |
| Diet 18 | 19.16 | 33.33 | 47.5 | 30.34 | 23.45 | 75.2 | 129.01 | 850.05 | 6.59:1 | 1:4.22 | 01:02 | 1:1.11 | 30.70 | 23.73 | 88.1 |
| Diet 19 | 19.16 | 47.5 | 33.33 | 30.34 | 33.82 | 52.77 | 116.55 | 866.25 | 7.43:1 | 1:4.22 | 1:1.11 | 01:02 | 30.70 | 33.81 | 61.83 |
| Diet 20 | 19.16 | 61.66 | 19.16 | 30.34 | 43.9 | 30.34 | 104.08 | 882.44 | 8.48:1 | 1.:4.22 | 1.61:1 | 1:4.22 | 30.70 | 43.90 | 35.55 |
| Diet 21 | 19.16 | 75.83 | 5.00 | 30.34 | 53.36 | 7.91 | 91.62 | 898.63 | 9.81:1 | 1:4.22 | 3.13:1 | 01:19 | 30.70 | 43.90 | 35.55 |
| **Diet 22** | **5.00** | **5.00** | **90.00** | **7.91** | **3.51** | **142.5** | **153.93** | **814.08** | **5.29:1** | **01:19** | **01:19** | **09:01** | **8.00** | **3.55** | **166.94** |
| **Diet 23** | **5.00** | **19.16** | **75.83** | **7.91** | **13.48** | **120.06** | **141.47** | **830.27** | **5.87:1** | **01:19** | **1:4.22** | **3.14:1** | **8.00** | **13.64** | **140.66** |
| Diet 24 | 5.00 | 33.33 | 61.66 | 7.91 | 23.45 | 97.63 | 129.01 | 847.47 | 6.56:1 | 01:19 | 01:02 | 1.61:1 | 8.00 | 23.73 | 114.38 |
| Diet 25 | 5.00 | 47.50 | 47.50 | 7.91 | 33.42 | 75.20 | **116.55** | **862.66** | **7.4:1** | 01:19 | 1:1.11 | 1:1.11 | 8.00 | 33.81 | 88.10 |
| Diet 26 | 5.00 | 61.66 | 33.33 | 7.91 | 43.39 | 52.77 | 104.08 | 878.85 | 8.44:1 | 01:19 | 1.61:1 | 01:02 | 8.00 | 43.90 | 61.83 |
| **Diet 27** | **5.00** | **75.83** | **19.16** | **7.91** | **53.36** | **30.34** | **91.62** | **895.05** | **9.77:1** | **01:19** | **3.14:1** | **1:4.22** | **8.00** | **53.98** | **35.55** |
| **Diet 28** | **5.00** | **90.00** | **5.00** | **7.91** | **63.33** | **7.91** | **79.16** | **911.24** | **11.51:1** | **01:19** | **09:01** | **01:19** | **8.00** | **64.07** | **9.27** |

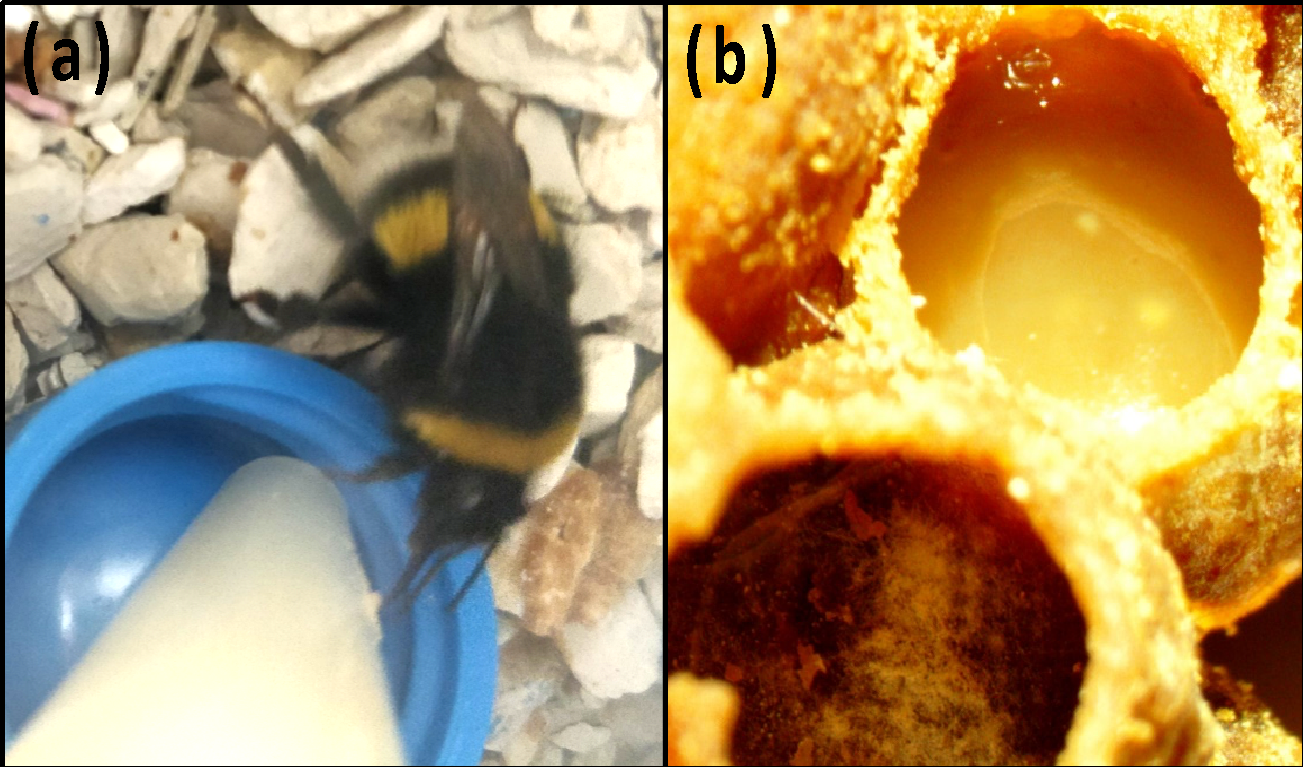

**Figure S1: Collection of artificial diet by bumblebees**. The artificial diets were provided in **(a)** gravity feeders (drilled Eppendorf of 5ml placed above a plastic cub), from which they were collected by bumblebees for immediate consumption or **(b)** storage in empty brood cells.

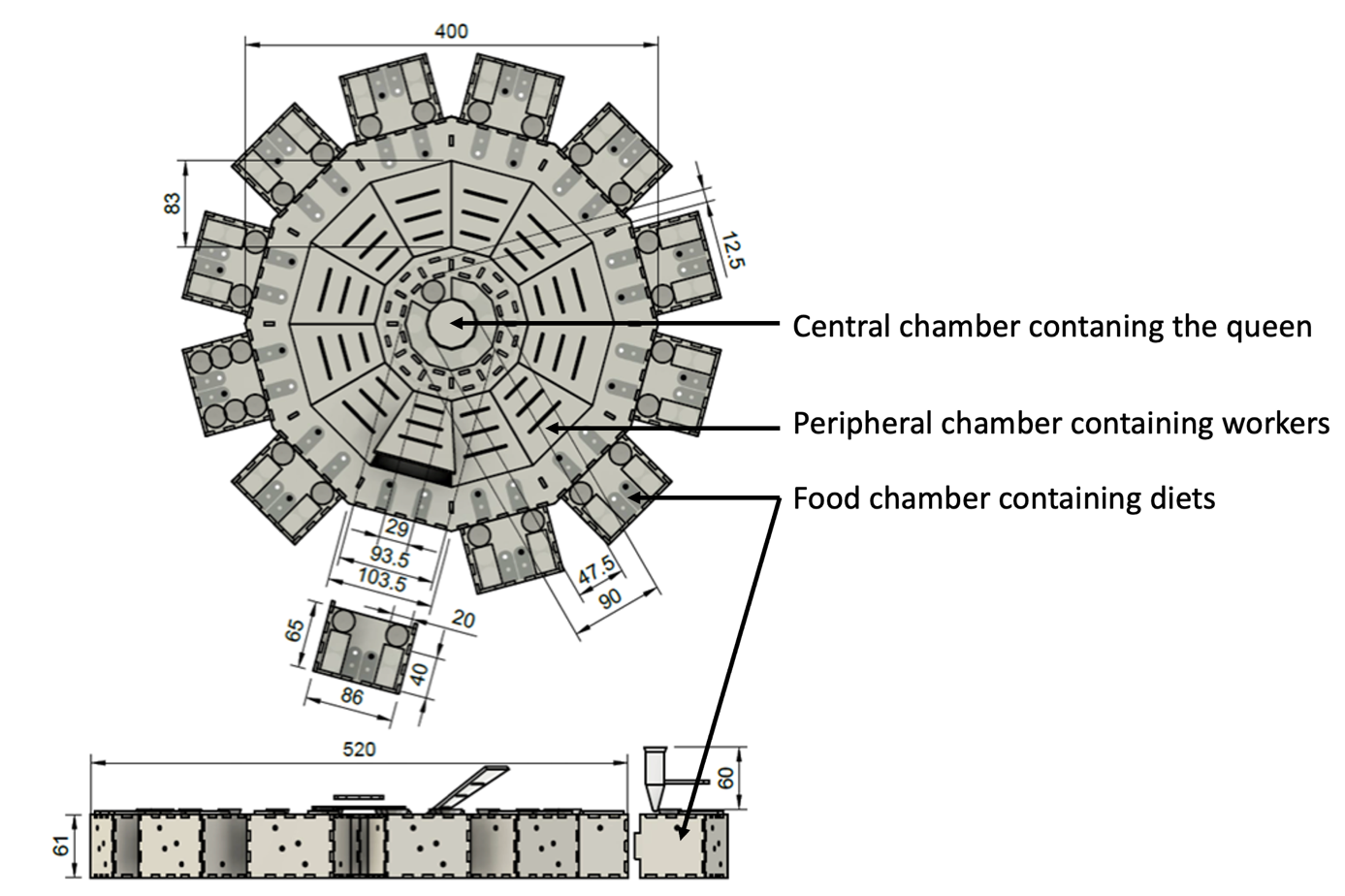

**Figure S2: Technical schematics of the dodecagonal arena used in the choice and no-choice experiments**. The arena (Ø520mm; height: 61mm) was made of a central chamber (Ø93.5mm; floor area: 6866mm²) surrounded by 12 peripheral chambers (length: 144mm; floor area: 9540mm²). The central chamber had two clear transparent walls with air gaps that precluded access to the workers’ chambers. The peripheral chambers were separated from each other by an opaque wall; each one gave access to a food chamber (86x65x61mm; floor area: 5590mm²) where 4 drilled Eppendorf vials of 5ml provided food. The ground was covered with cat litter. Central and peripheral chambers were covered with cardboard to protect from light exposure.

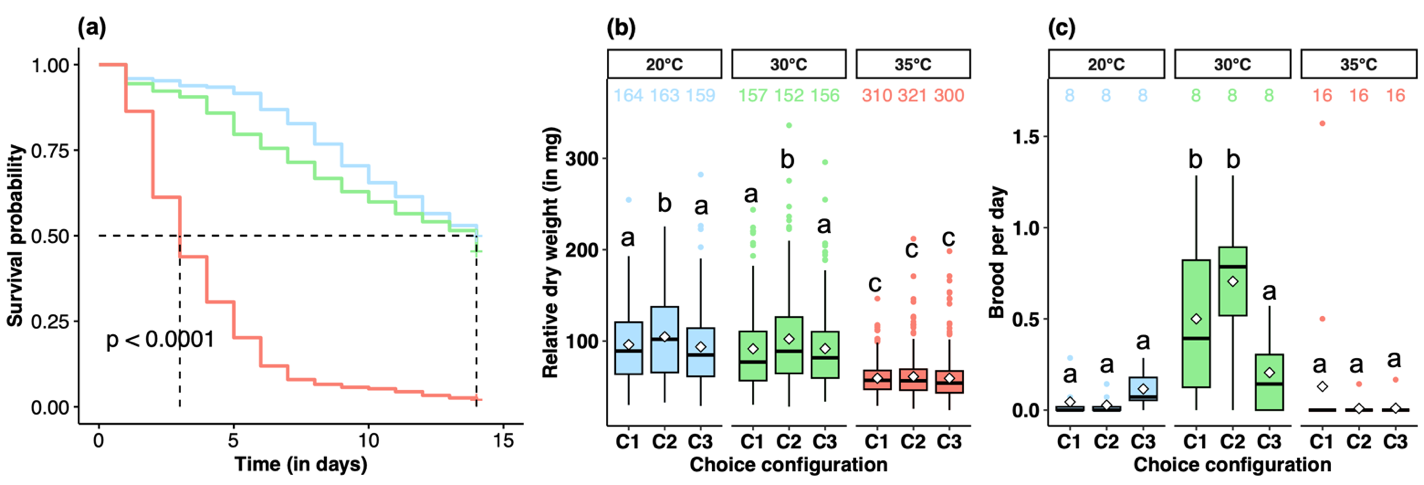

**Figure S3: Fitness performances for the three temperature regimes in the choice experiments**. **(a)** Survival in choice experiments with different temperatures. Survival curves from a Kaplan-Meier model, with confidence interval at 95% and the p-value. **(b)** Boxplots of relative dry weight (in mg) for a bee of median size (of 4.5mm thorax width). The central line is the median, the edges of the box are the 25th and 75th percentiles, the whiskers extend to the most extreme data points without considering outliers, which are represented by dots. Letters (a-c) show significant differences from GLMM.

**Text S1: Fitness performances in the choice experiments.** We measured survival and body weight for individual bumblebees during the 14 days of the choice experiments. The survival probability of the bees was not influenced by the interaction between choice configuration and temperature (ANOVA: Chi=7.591, df=4, p=0.108). The additive model showed no effect of the choice configuration (ANOVA: Chi=1.269, df=2, p=0.530), but an effect of the temperature (ANOVA: Chi=145.440, df=2, p<0.001). At 35°C, the bees had a 9.5 times greater probability of death than at 20°C or 30°C (Cox Model: estimate±standard error = 2.25±0.22, z = 10.21, p <0.001; Figure S3.a). The relative dry weight (in mg) for a median bee (of 4.5mm thorax width) was not influenced by as the interactions between choice configuration, temperature and mortality (ANOVA: Chi=4.776, df=4, p=0.311). However, we found a significant interaction between choice configuration and temperature (ANOVA: Chi=11.831, df=4, p=0.019) and between choice configuration and mortality (ANOVA: Chi=8.997, df=2, p=0.011). The bees that were still alive at the end of the experiment weighed 40.77 mg [-12;94.92] more than those found dead, as expected from a continuous intake during the experiment duration. In addition, bees were 1.5 times lighter at 35°C than at 20°C or 30°C (Figure S3.b).
